# Polarimetric response of second harmonic generation in microscopy of chiral fibrillar structures

**DOI:** 10.1101/2023.02.23.529792

**Authors:** Mehdi Alizadeh, Fayez Habach, Mykolas Maciulis, Lukas Kontenis, Saulius Bagdonas, Serguei Krouglov, Vytautas Baranauskas, Danute Bulotiene, Vitalijus Karabanovas, Ricardas Rotomskis, Margarete K. Akens, Virginijus Barzda

## Abstract

Polarimetric second harmonic generation (SHG) microscopy is employed to study partially oriented fibrillar structures. The polarimetric SHG parameters are influenced by three-dimensional (3D) configuration of C_6_ symmetry fibrilar structures in the focal volume (voxel) of a microscope. The achiral and chiral susceptibility tensor components ratios (R and C, respectively) are extracted from the linear polarization-in polarization-out (PIPO) measurements. The analytical derivations along with the polarimetric SHG microscopy results obtained from rat tail tendon, rabbit cornea, pig cartilage and meso-tetra (4-sulfonatophenyl) porphine (TPPS_4_) cylindrical aggregates demonstrate that SHG intensity is affected by parallel/antiparallel arrangements of the fibers, and R and C ratio values change by tilting the fibers out of image plane, as well as by crossing the fibers in 2D and 3D. The polarimetric microscopy results are consistent with the digital microscopy modeling of fibrillar structures. These results facilitate the interpretation of polarimetric SHG microscopy images in terms of 3D organization of fibrilar structures in each voxel of the samples.

**Statement of Significance:** Polarimetric second harmonic generation (SHG) microscopy is used to study partially oriented C_6_ symmetry chiral fibrillar structures. The linear polarization-in polarization-out (PIPO) SHG imaging is performed on rat tail tendon, rabbit cornea, pig cartilage tissues and meso-tetra (4-sulfonatophenyl) porphine (TPPS_4_) cylindrical aggregates. The study demonstrates that SHG intensity is affected by parallel/antiparallel arrangements of the fibers, and the achiral and chiral susceptibility component ratio values change by tilting the fibers out of image plane, as well as by crossing the fibers in 2D and 3D. The polarimetric microscopy results are consistent with the digital microscopy modeling of fibrillar structures. These results facilitate the interpretation of polarimetric SHG microscopy images in terms of 3D organization of fibrillar structures in each voxel of the samples.

## 1 Introduction

Polarimetric second harmonic generation (SHG) microscopy has been widely used for revealing the ultrastructure of biological specimens comprised of noncentrosymmetric chiral fibrilar structures. The fibrilar structures of collagen (1, 2), myosin (3, 4), and starch granules (5) that are partially oriented in the biological samples efficiently generate second harmonic signal, which is sensitive to polarization. The nonlinear polarimetric response can be employed to extract ultrastructural information about the molecular organization in each voxel of the imaged structures.

The ultrastructure of biological samples can be characterized using various polarimetric SHG microscopy techniques (6–12). Complex valued measurable nonlinear susceptibility tensor components for two-dimensional (2D) case can be extracted using complete double Stokes-Mueller polarimetry (10, 11, 13). The susceptibility component ratios and the effective fibrilar orientation in the image plane can be obtained with various reduced SHG polarimetry techniques by assuming structural constrains, i.e. C_6_ symmetry for fibrillar structures. By controlling the incident and outgoing linear polarization states, the chiral and achiral nonlinear susceptibility component ratios and the effective orientations can be deduced (7, 14). The achiral ratio and effective fiber orientation can also be extracted by employing only the incident linear polarizations (4, 9, 15, 16), or by using circular incident and linear outgoing polarization states (8, 12). The achiral susceptibility ratio can also be obtained with circular incident and outgoing polarization states in fast, fiber orientation independent measurements (17).

Many SHG active fibrillar molecules at various orientations can be present in a voxel since the diameter of constituting fibers is usually one order of magnitude smaller than the focal volume diameter of a high numerical aperture (NA) objective. Several configurations such as parallel/antiparallel fibers (18), tilted fibers out of image plane (14), and crossed fibers in 2D and 3D (14, 16, 19) significantly influence the SHG polarimetric response. Recently, a digital nonlinear microscope has been applied to model SHG polarimetric response of different fiber configurations in a voxel (20). In this article, we present characteristic SHG microscopy cases of fibrillar organizations appearing in biological and biomimetic samples that significantly influence the polarimetric response.

The study employs polarization-in polarization-out (PIPO) SHG microscopy (7, 14, 21) to investigate collagen ultrastructural organization in rat tail tendon, rabbit cornea and pig cartilage tissues, and chiral fibers of mesotetra (4-sulfonatophenyl) porphine (TPPS_4_) molecules in giant “sea urchin” (GSU) aggregates (22). Digital nonlinear microscopy modeling results published in the preceding paper (20) will be used to interpret the experimental polarimetric results. The study demonstrates that polarimetric SHG microscopy can reveal 3D ultrastructural organization of fibers within voxels of imaged specimens.

## 2 Theoretical background

### 2.1 PIPO equation for a chiral fibrillar structure

The linear PIPO SHG microscopy provides information on achiral R and chiral C susceptibility ratios that characterize the C_6_ symmetry fibrillar structure contained in a voxel. The SHG intensity relates to the incident and the outgoing linear polarization orientations as follows (7, 20):

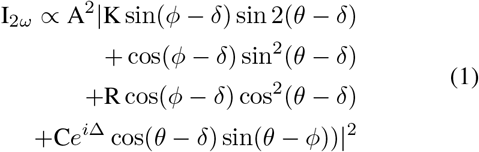

where *θ* and *ϕ* respectively are the incident and outgoing linear polarization orientation angles considered from the Z axis of the laboratory coordinate system. The *δ* is the orientation angle of cylindrical axis of a fibrillar structure measured from the Z axis. The sample plane is considered along XZ plane of laboratory coordinate system and Y is the beam propagation direction (Fig. 1). The amplitude A of the SHG response will depend on a configuration of fibers in a voxel, as will be defined later. The so called Kleinman ratio, K, is considered to be around K=1 for the off resonance conditions (7, 10). The R and C ratios depend on the organization of fibers in a voxel. In this paper, the main types of fiber organizations will be considered including a single fiber, and two fibers with crossing and parallel/antiparallel configurations.

**Figure 1:**
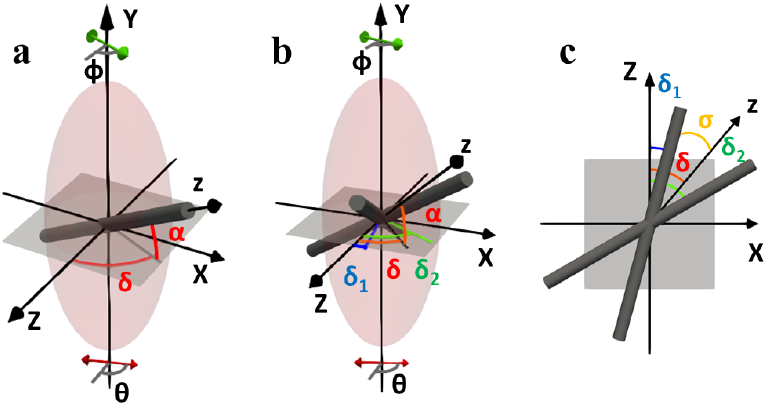
Laboratory coordinate system XYZ and molecular coordinate system xyz for cylindrical fibers with C_6_ symmetry in a voxel. (a) A single fiber in a voxel. The fiber is oriented with the in-image-plane (ZX) angle *δ* and the tilt angle *α*. The molecular z axis is along the fiber axis. (b) Two fibers in a voxel with individual angles *δ*_1_ and *δ*_2_, and the effective angle *δ* with the effective molecular axis z. The *α* angles are the same for both fibers. (c) The bottom view of crossing fibers projection on the image plane (ZX). *θ* and *ϕ* are the incoming and outgoing light polarization orientation angles, respectively, measured from the laboratory Z axis in (a, b). The light propagates along the laboratory Y axis.

### 2.2 A single fiber configuration in a focal volume

A single fiber may be present in a focal volume (Fig. 1a). The fiber is oriented with in-image-plane angle *δ* and out-of-image-plane tilt angle *α*. The molecular coordinate system has z axis along and x, y axes perpendicular to the fiber. The PIPO data can be fit with eq. (1) resulting in R, C and *δ* parameters. The amplitude A will be equal to the amplitude for one fiber A^*′*^, which is expressed as:

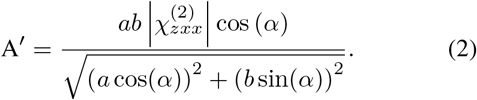

The *a* and *b* denote semi-major and semi-minor axial widths of the focal volume spheroid (see Fig. 1). 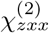 denotes the nonlinear susceptibility component in the molecular coordinate system.

For a single fiber, the Kleinman susceptibility ratio 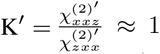 is assumed for off resonance conditions, where 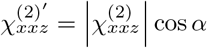 and 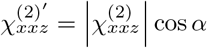

The achiral susceptibility ratio of a single fiber 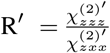 is assumed to be real valued, and can be expressed in terms of the susceptibility tensor elements in the molecular frame as follows (7):

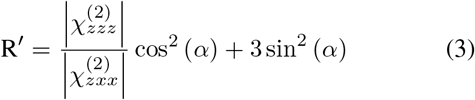

The chiral susceptibility ratio of a single fiber is considered to be complex valued with the magnitude 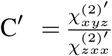 and the phase retardancy Δ between achiral and chiral susceptibility components (10, 23, 24):

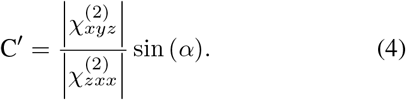

### 2.3 Two proximal fibers at variable crossing angles

Several fibers with different orientations may occur at a close proximity in the focal volume. Since the fibers are at a short distance from each other, the total far field SHG intensity can be calculated by simple addition of the amplitudes of electric fields generated from individual fibers, and the phase relations can be ignored (16, 20, 25). Here we outline the simplest case of two fibers with different in-image-plane orientations *δ*_1_ and *δ*_2_, and having the same *α* tilt angles (see Fig. 1b, c). This configuration represents two crossing fibers that preserves C_6_ symmetry of the whole structure. Therefore, PIPO equation (1) can be used for polarimetric data analysis. In this case, we define the effective in-image-plane orientation angle 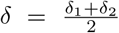 and the half crossing angle 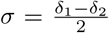 (see Fig. 1b, c). The primed susceptibility ratios of individual fibers R^*′*^ and C^*′*^ are considered to be the same, if *α* angles are the same for both fibers. For the PIPO data fit of crossed fibers with eq. (1), the Kleinman ratio K_*c*_ = 1 is assumed, and the fit parameters are:

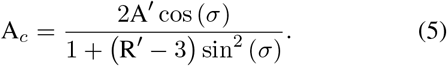

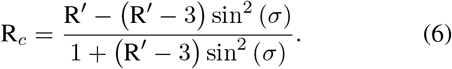

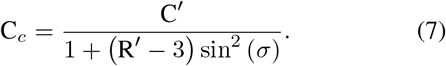

The dependence of R_*c*_ and C_*c*_ on the half crossing angle *σ* will be different if the R^*′*^ is below or above 3 (see Fig. 2). Note that R^*′*^ and C^*′*^ depend on the molecular susceptibility ratios 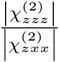 and 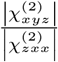, respectively, and the fiber tilt angle *α* according to eqs. (3, 4). The characteristic molecular values are taken 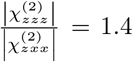 for collagen (7, 14, 18, 26, 27) and 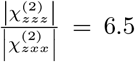 for TPPS (20, 22) fibers. For =0°, R′ will have the same value as the molecular susceptibility ratio (see eq. (3)). When tilted by *α*=45°, the fibers get R^*′*^=2.2 for collagen and R^*′*^=4.75 for TPPS_4_ according to eq. (3). The plots of eq. (6) in Fig. 2a demonstrate that R_*c*_ ratio increases for collagen and decreases for TPPS_4_ fibers by increasing the crossing angle 2*σ*. The R_*c*_ = 3 is reached for both collagen and TPPS_4_ aggregates when the crossing angle assumes 60°(see eq. (6)). R_*c*_ = 3 also appears at the critical value of 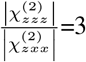 and does not depend on *α* or *σ*.

**Figure 2:**
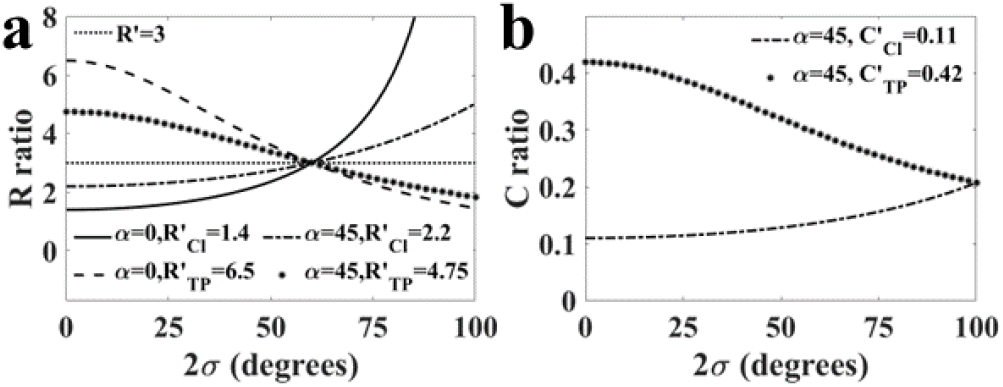
The effective R_*c*_ (a) and C_*c*_ (b) ratio dependence on the crossing angle 2*σ*=(*δ*_1_ − *δ*_2_) between the two fibers with the molecular 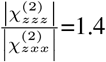 for collagen, 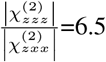 for TPPS_4_ aggregates, and 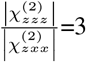 for critical fibers. 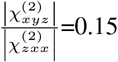 for collagen and 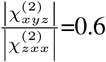 for TPPS_4_ fibers is considered. 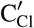 for collagen and 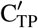 for TPPS_4_ are calculated from eq. (3) with *α* tilt angle of 0°or 45°, see figure legend. Correspondingly, 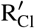 and 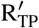 are calculated from eq. (4) for positive *α*.

For the crossing fibers, C_*c*_ ratio depends on C^*′*^, but also on R^*′*^ and half crossing angle *σ* (eq. (7)) as shown in Fig. 2b. C_*c*_=0 if fibers are in the image plane. When fibers are tilted out of the image plane C^*′*^ acquires some value, and C_*c*_ value increases with the increase of crossing angle 2*σ* for collagen and decreases for TPPS_4_ fibers (Fig. 2b). The effect is opposite for the negative C^*′*^ fibers. When R^*′*^=3, the C_*c*_=C^*′*^ and C_*c*_ does not depend on the half crossing angle *σ*. The simulation results of crossing fibers (Fig. 2) are consistent with the modeling of different fiber configurations in the focal volume obtained by the digital microscope (20).

### 2.4 Parallel and antiparallel fibers

The fitting parameters of PIPO equation for the parallel and antiparallel fiber configurations can be expressed from the crossing fibers equations (5), (6) and (7). For two parallel fibers *σ* = 0°, which gives A_*p*_ = 2A^*′*^, R_*p*_ = R^*′*^ and C_*p*_ = C^*′*^. For the antiparallel fibers *σ* = 90°, and A_*a*_ = 0, which indicates that SHG intensity vanishes and R_*a*_ and C_*a*_ can not be retrieved in the experiment.

## 3 Materials and methods

### 3.1 PIPO microscope

The PIPO microscope setup has been fully described previously (7, 14, 28–30). Briefly, two home built microscopy setups were used with minor differences between them. For imaging the pig cartilage and TPPS_4_ aggregates, the custom-designed diode-pumped Yb-ion-doped potassium gadolinium tungstate (Yb:KGW) crystal-based oscillator (31) was used. The laser generates 430 fs pulses at 14.3 MHz repetition rate and operates at 1028 nm wavelength. Rat tail tendon and rabbit cornea samples were imaged with the second microscope using pulsed femtosecond oscillator (FLINT, Light Conversion, Lithuania) that generates 100 fs pulses with 76 MHz repetition rate at 1030 nm central wavelength. The lasers were coupled to the respective nonlinear microscopes. The sample scanning was accomplished with a pair of galvanometric mirrors. For excitation in the first microscope, a 20 × 0.75 NA air objective (Carl Zeiss) was used and the signal was collected with a custom 0.8 NA objective. For the second microscope, 20 × 0.75 NA air objective (Nikon Plan Apo Lambda) was used and the signal was collected with a singlet 0.45 NA lens. The SHG signal was detected by a single-photon counting PMT (Hamamatsu H7422P-40, for the first microscope, and H10682-210, for the second). To separate SHG from the laser radiation a BG 39 Schott color glass filter and a 510 nm to 520 nm wavelength band-pass interference filter were used. A polarization state generator (PSG) containing a linear polarizer and a half-wave plate was placed before the excitation objective. A polarization state analyzer (PSA) was placed after the collection lens and had a rotating linear polarizer. In order to perform PIPO measurements, the PSG half-wave plate was rotated from 0°to 180°in steps of 22.5°, and the PSA was rotated in the same way as PSG from 0°to 180°for each PSG state. Also, to monitor sample photostability, the samples were rescanned at the initial PSG and PSA settings after finishing the measurements with each set of the PSA states.

### 3.2 Sample preparation

Rat tail tendon (RTT) samples were prepared from 12 weeks old Albino Wistar rats acquired from the State Scientific Research Institute of Innovative Medical Center (Vilnius, Lithuania). The collected tendon tissue was sectioned to 5 *μm* slices at longitudinal and oblique orientations with respect to the collagen fibers using a cryostat microtome. The sections were mounted on microscope slides and stained using a standard haematoxylin and eosin (H&E) staining procedure. All animal procedures were carried out in accordance with the guidelines set by the State Food and Veterinary Service Animal Care and Use Committee (Vilnius, Lithuania), which approved the current study (approval number G2-156).

Rabbit corneas were dissected following euthanasia. Tissues were fixed overnight in 4% paraformaldehyde and sectioned with a cryostat microtome. The tissue sections of 8 *μm* thick were stained by a standard H&E staining procedure. Animals were treated in accordance with the Association for Research in Vision and Ophthalmology guidelines for the use of animals in ophthalmic and vision research, and the directive 2010/63/EU on protecting animals used for scientific purposes, using protocols approved and monitored by the State Food and Veterinary Service, Lithuania, Vilnius University animal license number G2-95.

Juvenile swine tissue was harvested from a 25 kg Yorkshire pig euthanized within another approved research study at the University Health Network, Toronto, Canada. The articular cartilage tissue was fixed in 10% buffered formalin and cut perpendicular to the cartilage surface. The samples were embedded in paraffin, and sectioned into 5 *μm* thick slices. The slices were deparaffinated, mounted on microscope slides and stained with hematoxylin and eosin using standard procedures.

The filaments of meso-tetra(4-sulfonatophenyl)porphine (TPPS_4_) aggregates from the giant sea urchin (GSU) type aggregate preparations were used for investigations (22). The preparation of TPPS_4_ aggregates is described elsewhere (22). Briefly, a 10^−3^ M stock solution of TPPS_4_ was prepared by dissolving the powder (Sigma Aldrich, St. Louis, Missouri, USA) in distilled water. The fibrillar aggregates and GSU supramolecular structures self-assembled by diluting the stock to 2 × 10^−5^ M with 0.1 M aqueous HCl solution under random mixing. The solution was aged in tightly closed 50 ml volume flasks in the dark at room temperature for more than a year. The TPPS_4_ aggregates were mounted in the original aggregation medium on the microscope slides and covered with coverslips for imaging.

## 4 Results

### 4.1 Parallel and antiparallel collagen fibers in a tendon

Tendon tissue structure is comprised of parallel and antiparallel fibers (6) resulting in stripy appearance of the tissue in SHG intensity images shown in Fig. 3b, g (32, 33). In contrast, the multiphoton excitation fluorescence (MPF) of the same scan areas exhibits almost homogeneous intensity images (Fig. 3a, f). MPF originates from eosin that stains proteins including collagen in H&E stained tissue (34). The uniform appearance of tendon tissue in MPF images indicates that stripy structures observed in SHG intensity images are likely due to coherent signal summation of varying distribution of parallel and antiparallel fibres (10, 18, 19). This phenomenon is predicted by numerical modeling of antiparallel fibers with the digital nonlinear microscope (20), and also described in the theory section for antiparallel fibers.

**Figure 3:**
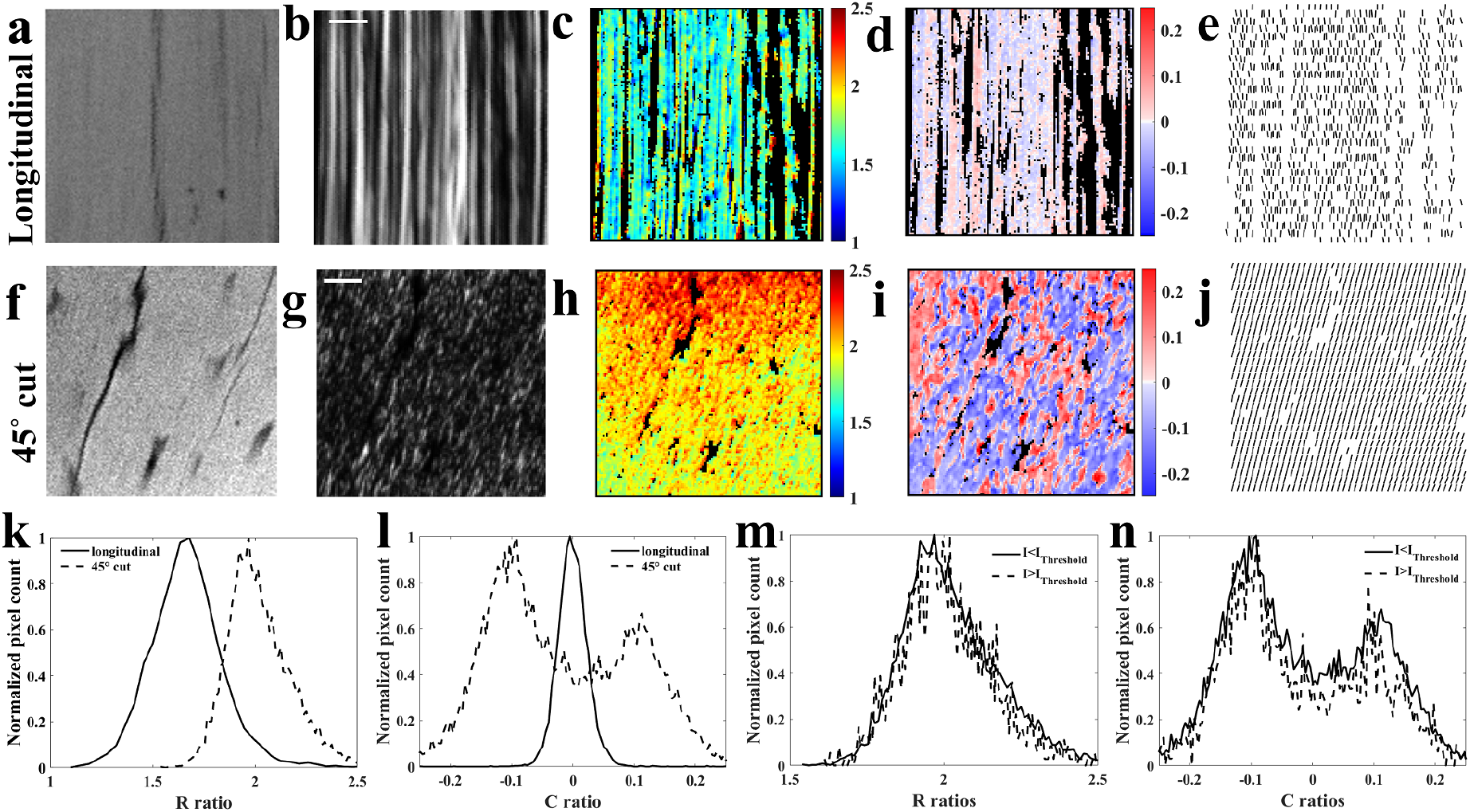
PIPO microscopy of rat tail tendon collagen for longitudinal (a-d) and 45°(f-j) cut. MPF images of the two samples (a and f). SHG intensity (b and g) obtained by summing all PIPO images of the corresponding samples. R ratio maps (c and h) calculated from the corresponding PIPO images. C ratio maps (d and i). In-image-plane effective orientation *δ* maps (e and j). Normalized histograms of R ratio (k) for samples corresponding to (c and h) ratio maps. Normalized histograms of C ratios corresponding to (d and i) ratio maps. Normalized histograms of R ratios (m) and C ratios (n) of the pixels below (solid line) and above (dashed line) 30% threshold of the SHG intensity for 45°cut sample. Scale bar is 5*μm*.

The PIPO results of the imaged areas provide with the effective R ratio maps (Fig. 3c, h). A relatively narrow R ratio distribution for the longitudinal cut tendon peaks at R_mode_ ≈ 1.67 (Fig. 3k), which is in a good agreement with the reported R ratios for rat tail tendon (35). A much larger R ratio values appear when the fibers are tilted out of the image plane (Fig. 3h) (14). The mode of R ratio histogram increases to R_mode_ = 2 for 45°cut. This effect is visible by comparing the normalized histograms of R ratios for the longitudinal and 45°cut tendon (Fig. 3k). The observed R ratio dependence on tilt is in a good agreement with the results predicted by eq. (3) and the numerical modeling of the digital PIPO microscope (20). The R ratio map of the 45°cut tendon (Fig. 3h) shows higher values at the upper part and lower values at the bottom of the image. This occurs due to differences in tilt angle *α* across the area. It can be seen from SHG intensity image (Fig. 3g) that longer streaks corresponding to the fibers cut at shallower *α* angle appear in the lower part of the image compared to the upper part region with dot like appearance of the fibers corresponding to larger *α* tilt angles. Thus, visual variation of tilt angle *α* correlates well with R ratio distribution across the image in Fig. 3h.

The corresponding C ratio maps are shown in Fig. 3d and i. The reddish areas correspond to positive tilt angles out of the image plane and the blueish areas show negative tilts. The small C ratio values for the longitudinal cut indicate that most of the fibres are oriented predominantly in the image plane (Fig. 3d). The C ratio values are larger in the tilted case and both negative and positive (blue and red) values can be observed in the image Fig. 3i, showing varying polarity organization of the fibers in tendon tissue. The normalized histograms of C ratios are shown in Fig. 3l. The narrow peak of C_mode_ = 0 confirms that the longitudinal fibers are in the image plane. In contrast, a broad distribution of C ratio with two peaks (positive and negative) for the 45°cut fibers shows the presence of opposite polarity fibrillar organization in the tendon. The results match numerical modeling of antiparallel fiber configurations for collagen (20).

The SHG intensity pixel values do not correlate with the corresponding R and C ratio values (compare panels g with h and i). The stripes with high SHG intensity (above 30% threshold of the maximum) have the same distributions of R and C ratios compared to the low SHG intensity pixels (below 30% threshold), as shown by the normalized R and C ratio histograms for the 45°cut in Fig. 3m and n, respectively. This is in line with the modeling where different amount of parallel/antiparallel fibers results in a large variation of intensity due to coherent summation of SHG signal, while similar values of R and C ratios are obtained (20).

Fig. 3e and j show the in plane effective orientation maps of fibers in the corresponding areas, which confirm the parallel (antiparallel) orientation of fibers in the tendon tissue. Therefore, stripy appearance of the tendon tissue in SHG intensity images and island like distributed positive and negative values of C ratio maps at 45°tendon cut point to the parallel/antiparallel arrangement of the collagen fibers in tendon tissue.

### 4.2 2D crossing fibers in rabbit cornea

Cornea is composed of stacked multiple lamellae of type I collagen fibers. Due to different orientations of collagen in the corneal tissue 2D crossing of fibers can be observed with SHG microscopy (16, 36–38). Longitudinal cuts of normal rabbit corneas have been investigated with PIPO SHG microscopy (Fig. 4). SHG intensity maps of non-crossing and crossing fiber areas are shown in Fig. 4a and e, respectively. Since PIPO polarimetry measurement sequence uses linear incident and linear outgoing polarization states at different orientations, when the polarization states are aligned with the fiber axis, the maximum SHG intensity is observed. Therefore, individual fibers can be highlighted at different orientations in the crossing area. Fig. 4k and l show two images corresponding to the PSG and PSA orientations at *θ* = *ϕ* =22.5°and *θ* = *ϕ* =112.5°, respectively. Comparison of the blue and green framed areas in Fig. 4k and l shows that blue frame area contains almost perpendicular crossing of two layers of fibers, while green frame area has predominantly unidirectionally oriented fibers at 112.5°from the vertical axis. SHG intensities of the crossing fiber areas are similar to the non-crossing areas.

**Figure 4:**
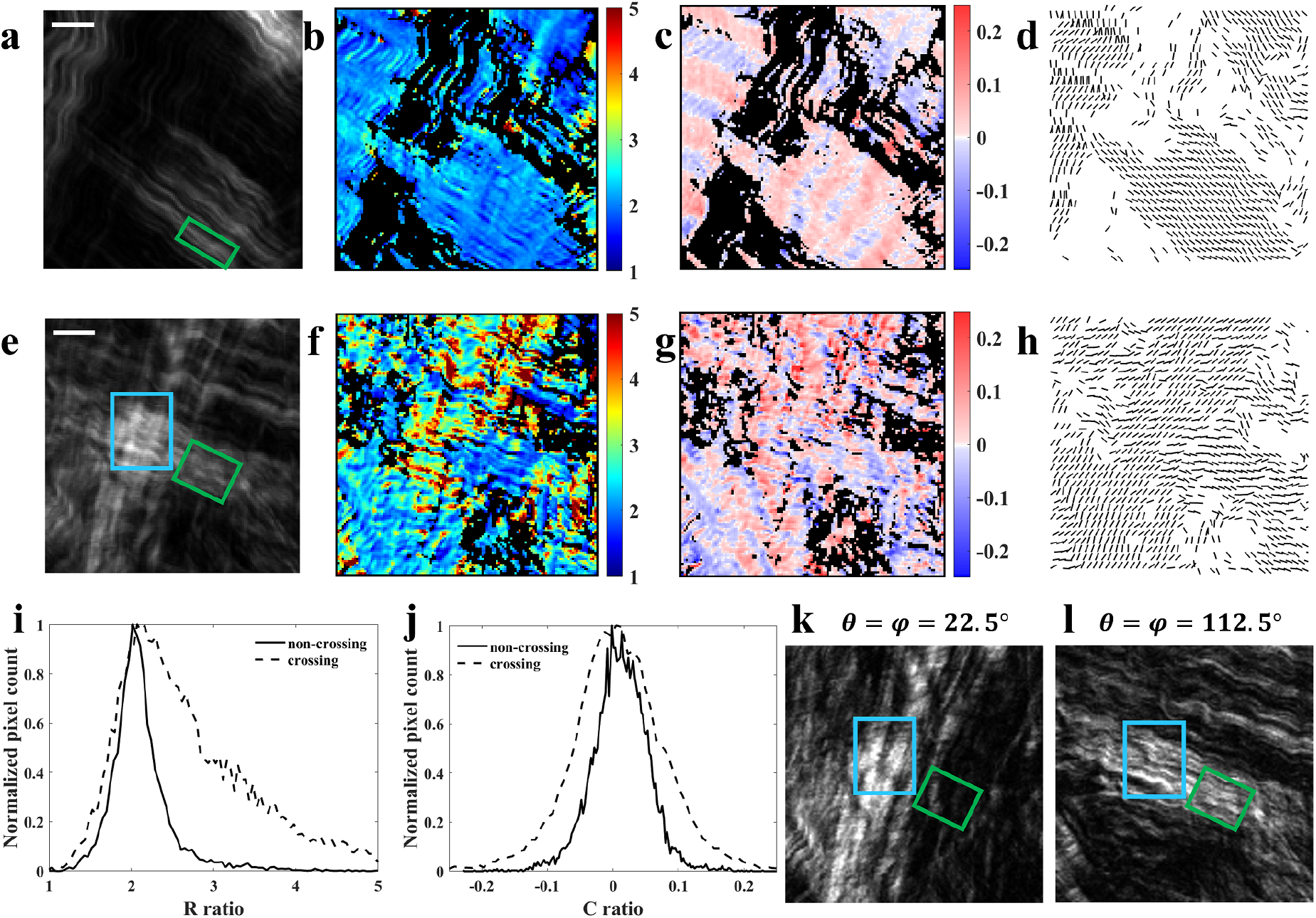
PIPO SHG microscopy of cornea containing non-crossing (a-d) and crossing (e-h) collagen fibers. SHG intensity (a, e), R ratio maps (b, f), C ratio maps (c, g) and in-image-plane effective fiber orientation maps (d, h) are shown. Normalized R and C ratio histograms are given for non-crossing (solid line) and crossing (dashed line) areas (i) and (j), respectively. SHG intensity of crossing area corresponding to (a) are presented for the polarization states with *θ* = *ϕ* =22.5°(k) and *θ* = *ϕ* =112.5°(l). The scalebar is 5*μm* in (a) and (e).

Crossing fibers have a strong effect on the R ratio, especially close to the 90°crossing angle (20). The R ratio maps calculated from non-crossing fiber areas have the ratio values close to R = 2 (see Fig. 4b), while crossing fiber areas show R values between 2 and 5 (Fig. 4f). The normalized histograms of R ratios (Fig. 4 i) for non-crossing (solid curve) and crossing (dashed curve) areas obtained from Fig. 4b and f show a narrow peak at R_mode_ = 2 for the non-crossing area and a wide peak at R_mode_ = 2.1 for the crossing area with higher values from 2 to 5. The presence of higher values indicates fiber crossing at large angles in the crossing area. If we take the half crossing angle to be *σ* = 45°and a single fiber R^*′*^ = 2, the calculated ratio of crossing fibers according to equation (6) is R_*c*_ = 5, which is observed in the crossing area of the sample.

The C ratio images show characteristic crimping structure of collagen fibers with up and down tilt wavy appearance resembling woven fabric in the crossing area Fig. 4c, g. The magnitude of C ratio values appears slightly larger for the crossing fibers. The normalized C ratio histograms of the non-crossing and the crossing areas are shown in Fig. 4j by solid and dashed lines, respectively. The width of the C ratio histogram for the crossing area is slightly larger. The wider distribution for crossing area is in line with equation (7). In addition, larger C ratio appears due to the fact that in crossing areas collagen fibers pass above/below one another instead of intersecting physically. Therefore, the fibers are slightly tilted up and then down in the crossing areas. Therefore, the histogram of C ratio values gets symmetrically broadened due to crossing and tilting of the fibers.

The orientation maps show that most fibers are arranged parallel in the layers (Fig. 4d and h) and the fiber crossing is almost perpendicular in the crossing area of the sample. The effective fiber orientations appear half way between the directions of the crossing fibers, which is reflected in the bar orientations of the crossing points in the panel Fig. 4h.

The lowest values for R ratio can be found in the areas where the C ratio is low and the fibers have unidirectional orientation. The analytical derivations (Fig. 2) and digital microscope modeling show that R ratio reaches above 3 when the crossing angle is higher than 60°. The highest values for R ratio in the PIPO measurements appear when the fibers are crossed and tilted out of the image plane i.e. showing also high C ratio (Fig. 4f, g). This effect is in good agreement with PIPO digital microscope modeling for simultaneously tilted and crossed collagen fibers (20).

### 4.3 3D crossing collagen fibers in cartilage

Cartilage tissue is composed of 3D crossing collagen fibers (39–41). The organization of fiber crossing depends on the depth from the surface of articular cartilage, and affects not only the mechanical properties (42–44), but also the nonlinear optical polarimetry parameters of the tissue (35, 45–47). Sections of cartilage tissue cut perpendicular to the surface are used for investigation of depth dependent organization of 3D crossing fibers. Fig. 5 shows the results of PIPO SHG microscopy measurements close to the surface of articular cartilage at depth of ≈ 100*μm* (panels b-e) and at depth of ≈ 800*μm* (panels f-i, see scan areas in panel a). Fig. 5b and f present logarithmic scale SHG intensity images summed over all PIPO polarization states. The images reveal crossing fiber organization at the two depths. SHG intensity images of selected polarization states are shown with *θ* = *ϕ* =0°in panel l and with *θ* = *ϕ* =90°in panel m to better reveal the organization of crossing fibers. The highlighted collagen fiber orientations change at different polarization orientations revealing the fiber crossing organization.

**Figure 5:**
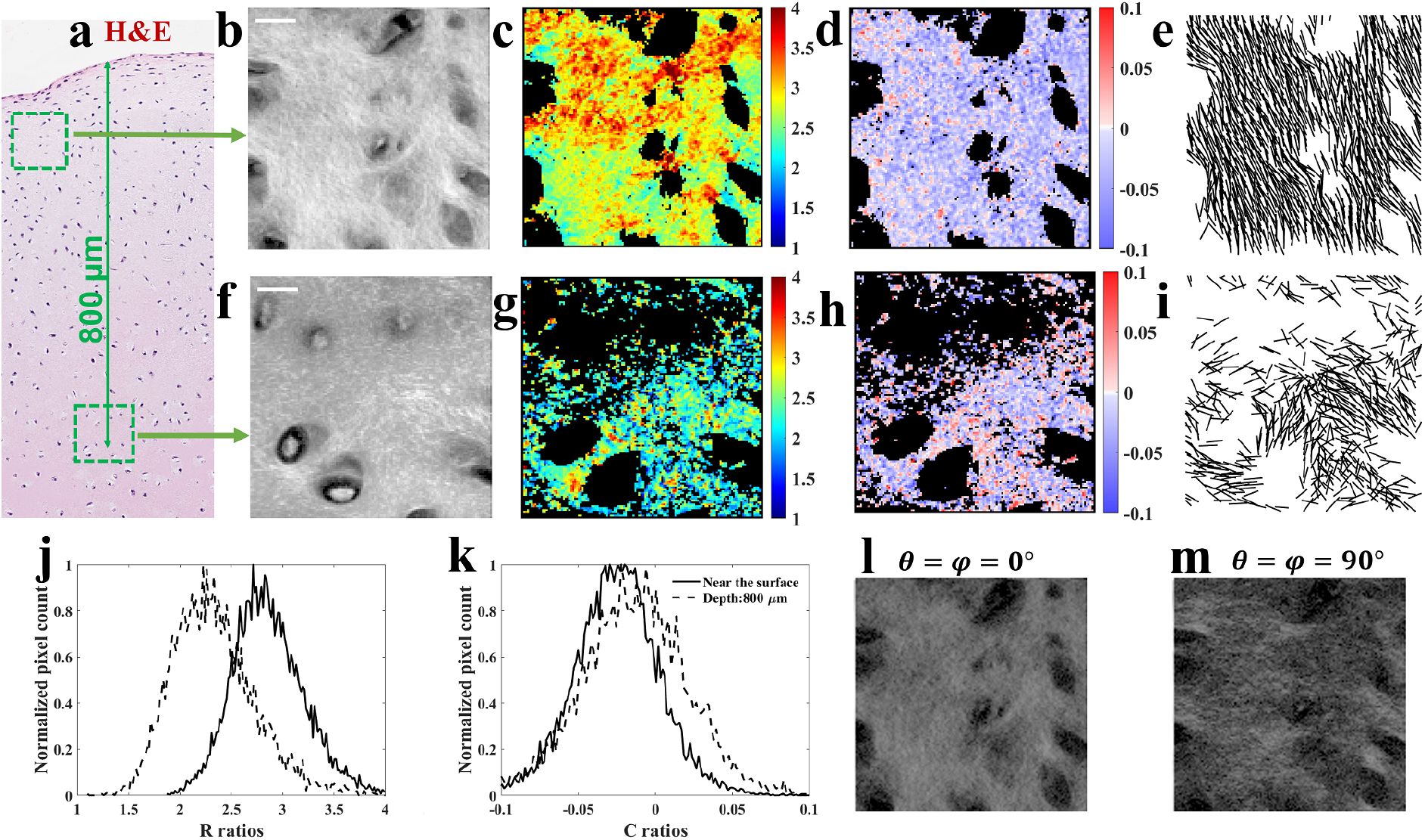
PIPO SHG microscopy of articular cartilage sectioned perpendicular to the cartilage surface. The polarimetric imaging is performed near the surface (b-e) and at the depth of 800*μm* (f-i). The corresponding scan areas are shown in H&E white light image (a). The polarimetric data contains logarithmic SHG intensity images (b, f), and achiral susceptibility R ratio (c, g), chiral C ratio (d, h) and effective fiber orientation (e, i) maps. The normalized histograms of R (j) and C (k) ratio values are presented for the areas near the surface (solid lines) and deeper in the tissue (dashed lines). SHG intensity images of the area near cartilage surface are presented for *θ* = *ϕ* =0°(l) and *θ* = *ϕ* =90°(m) polarization states. Scale bar is 20*μm* in (b, f).

Fig. 5c and g show the corresponding maps of R values obtained with PIPO measurements. R values depend on the depth from the surface showing higher values at the surface of the articular cartilage. This effect has been investigated in four juvenile swine cartilage samples and ten areas were scanned. In all areas the R ratio increases closer to the cartilage surface. The R value histograms (Fig. 5j) clearly demonstrate a shift to the higher values for the area near the surface. The mode of R value distributions decreases from R_mode_ = 2.71 for the area close to the surface to R_mode_ = 2.23 at the depth of 800*μm*. There is also a significant number of pixels with R values above 3, which indicates the presence of crossing fibers according to equation (6) (see Fig. 2). There are several reasons affecting the R ratio. The change in R value with depth may occur due to the change in the molecular structure i.e. type of collagen at different depths of the cartilage. Another reason might be because of the difference in 3D crossing angle of fibers at different depths from the surface of articular cartilage. The PIPO digital microscope results show that for the collagen fibers organized in a cone structure the R ratio increases by increasing the fiber crossing angle in the cone (20).

The C ratio maps are depicted in Fig. 5d and h. Positive and negative C values appear randomly distributed across the area showing varying out-of-image-plane tilt angle of different polarity collagen fibers in the tissue. Slightly broader distribution of C ratios is observed at the larger depth (see panel h). The corresponding histograms for C ratio are plotted in Fig. 5k. The standard deviation of the C ratio distribution at the area near the surface is 0.0281 and at the depth of 800*μm* is 0.0356 indicating slightly larger spread of fiber orientations at larger depth. This result is in line with larger spread of in-image-plane orientation angles *δ* for larger depth visible in the effective orientation map of Fig. 5i, compared to the panel e of scanned area at shallower depth.

The depth dependent R, C ratios and the in-image-plane orientation angle spread for cartilage collagen can be interpreted with hierarchical structure of 3D crossing fibers, where the larger cone crossing angle appears in voxels closer to the surface of the articular cartilage. The effective cone axis is oriented predominantly perpendicular to the articular cartilage surface, and the orientation spread of the effective cone axes across the scanned area increases going deeper into the cartilage. Alternatively, the changes in the R, C values with depth might appear due to the change in concentration of different collagen types having different molecular susceptibilities of individual collagen fibers (35, 48).

### 4.4 Non-crossing and crossing fibers of TPPS_4_ aggregates

The fibrillar TPPS_4_ aggregates generate strong second harmonic signal and can serve as model structures having large molecular R ratio (22). High R ratios are observed, for example, in starch granules (49, 50). Fig. 6 shows two areas of the sample one with predominantly non-crossing (upper row) and another with crossing (lower row) fibers. The logarithmic SHG intensity images are presented in Fig. 6a and e and the R ratio maps are shown in Fig. 6b and f for non-crossing and crossing areas, respectively. The R ratio map has higher values in the non-crossing area, while the crossing area has lower R values. The histograms of R ratios have been compared in Fig. 6i. As predicted by the PIPO digital microscope (20) and analytical derivation (see Fig. 2), the crossing fiber configurations decrease R ratio for TPPS_4_ aggregates since the fibers have molecular achiral susceptibility ratio above 3. The mode of R ratio distribution is (R_mode_ = 5.07) for the non-crossing area, while (R_mode_ = 3.07) is observed for the crossing area. This is a large change in R ratio. The values of R around 3 indicate a broad spread of orientations of the crossing fibers.

**Figure 6:**
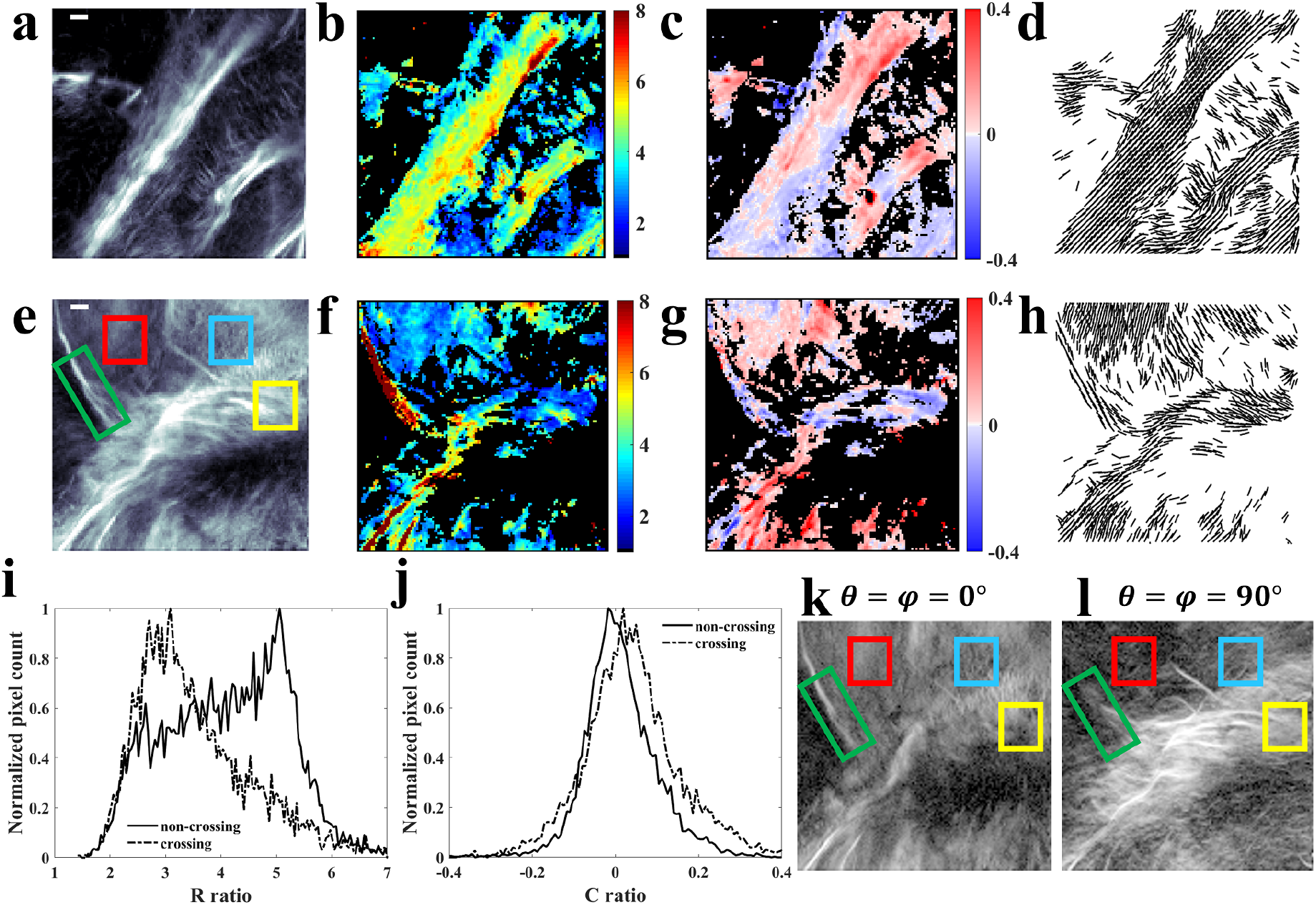
Samples of GSU TPPS_4_ aggregates predominantly without (a-d) and with (e-f) crossing fibers. Logarithmic SHG intensity images (a, e), achiral susceptibility R ratio maps (b, f), chiral susceptibility C ratio maps (c, g), and *δ* angle fiber orientation maps (d, h) are shown. Histograms of R ratio (i), and C ratio (j) values are given for non-crossing (b, c) and crossing (f, g) areas. Logarithmic SHG intensity frames are shown for *θ* = *ϕ* =0°(k) and *θ* = *ϕ* =90°(l) polarization orientations from the vertical frame axis. The indicated areas in (e, k) and (l) in green frame represent noncrossing, in red - crossing, in yellow - crossing and tilted, and in blue - broad orientation distribution of crossing fibers. Scale bar corresponds to 10*μm* in (a, e).

Fig. 6c and g show the C ratio maps. The histograms of C ratios are compared in Fig. 6j. The histograms have symmetric distributions around 0 value. The mode and the width of C ratio distributions are similar for both areas. Similar widths of C ratio histograms for crossing and non-crossing fibers suggest similar distributions of tilt angles *α* and/or chirality of the fibers in both samples (see Fig. 6j). Fig. 6d and h show the fiber orientation maps. The orientation map for non-crossing area contains mostly parallel bars, while crossing areas contain bars at different orientations. The crossing areas have correspondingly lower R ratios.

The crossing area image Fig. 6e includes several subareas that are interesting to highlight. There is a fiber inside the green highlighted area that presents the highest R ratio. By looking at different intensity images captured from different polarization states (see Fig. 6k and l) one can find out that this fiber has a unidirectional orientation without other fiber crossings. Fig. 6k shows that when both incoming and outgoing linear polarization orientations are aligned with 0°the fiber barely can be seen, yet it appears intense when the linear polarization orientations are at 90°with respect to the laboratory frame of reference (see Fig. 6l). Furthermore, both SHG intensity and R ratios are high in this area, consistent with the prediction of digital PIPO microscope modeling results of parallel fibers having R ratio around 6.5.

The effect of crossing fibers can be observed in the area inside the red square. The R ratio decreases dramatically in this area. The C ratio is almost zero indicating that majority of the fibers are in the image plane. R ratio reaches the lowest values in crossing area indicated with the yellow square, which contains also high C ratio values indicative of fibers tilted out of the image plane. The modeling with digital PIPO SHG microscope shows large R ratio decrease for the crossing and tilted fibers (20). The blue area highlights almost isotropically distributed crossing fibers. The area has weak SHG signal similar to Hyper-Rayleigh scattering (51), and the PIPO model for C_6_ symmetry cannot retrieve R and C ratios in this area. The presented PIPO SHG investigation for TPPS_4_ fibers shows that R ratio is very sensitive to crossing fiber configurations when the molecular R ratio values are large than 3.

## 5 Discussion

The impact of different fiber configurations on the polarimetric response of PIPO SHG microscopy measurements has been investigated using rat tail tendon, rabbit cornea, pig articular cartilage, and fibrillar TPPS_4_ aggregates. Three main fiber configuration types can be distinguished that largely influence the polarimetric SHG response: (i) the parallel/antiparallel fiber configuration, (ii) tilted fibers out of image plane, and (iii) crossed fibers in 2D and 3D. The combination of all three effects often is observed in biological tissue. Below, each fiber configuration effect is discussed separately.

The numerical modelling of PIPO digital microscope indicates that SHG intensity dramatically increases in the far field for parallel fibers and diminishes for antiparallel configuration due to coherent summation of the signal (20). The parallel/antiparallel organization of the fibers results in a stripy appearance of collagen in biological tissues, in contrast to the homogeneous appearance of connective tissue in eosin fluorescence image of H&E stained sample (Fig. 3 a), indicating that collagen concentration remains similar over the imaged area. This is in line with the previous studies on collagen polarity (33, 52, 53). R and C ratios are largely unaffected by the variation in SHG intensity of parallel/antiparallel fiber structures (Fig. 3 m, n) (20).

Tilting of the fibers out of image plane has a significant impact on the polarimetric nonlinear microscopy results (7, 14). R ratio increases with *α* increase for the structures with molecular susceptibility ratio 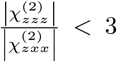 and decreases for the 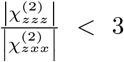. The molecular examples for 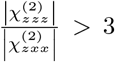 are collagen fibers with 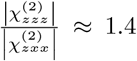 and for 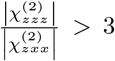 are TPPS_4_ aggregates with 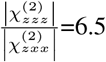. The imaging results demonstrate that R ratio increases with tilt for the tendon sample when comparing the longitudinal and the oblique (45°) tissue cuts in Fig. 3. The opposite effect of R is observed with tilt for TPPS_4_ aggregates (Fig. 6). The effect of tilt angle on C ratio is visible from the longitudinal and oblique cut tendon tissue images (Fig. 3). The C ratio distribution is centered around 0 for the longitudinal cut and becomes bimodal distributed with positive and negative peaks for the oblique cut due to opposite polarity of the fibers correspondingly tilted by *α* and − *α* angles (Fig. 3 l). The corresponding effect with tilt is observed for TPPS_4_ fibers in yellow framed area in Fig. 6, where R ratio is decreased while C ratio is increased compared to the green framed area with high R and low C ratio values containing in-image-plane oriented noncrossing fibers.

The crossing fibers influence R and C ratios. Since PIPO microscopy uses different polarization states, it is convenient to check for the fiber crossing by looking at the SHG intensity images taken with different linear polarization orientations. Each of the crossing fibers is highlighted when the linear polarization orientation matches individually the orientation of each fiber (R^*′*^ *>*1). For the structures with R^*′*^ *<*3, the R ratio increases with increasing the half crossing angle *σ*, and decreases for the structures with R^*′*^ *>*3. These results have been demonstrated using samples of collagen in rabbit cornea (Fig. 4) and TPPS_4_ aggregates (Fig. 6, red framed area in e, k, l). Similarly, the effective C increases with the crossing angle *σ* for R^*′*^ *<*3 and decreases for R^*′*^ *>*3, if the crossing fibers are tilted out of the image plane. 3D crossing fibers also appear in biological structures. The samples of articular cartilage are used to investigate the effect of 3D crossing collagen fibers on SHG polarimetric parameters. The depth dependency of effective R ratio is clearly observed in the cartilage sample. It might be due to the change in 3D crossing configuration of collagen fibers with depth. Alternatively, the depth dependent content of different collagen types in the cartilage may influence the changes in R ratio.

Different fiber organizations are present in biological tissues. These organizations can be decomposed into several basic fiber configurations affecting polarimetric SHG microscopy results, namely parallel/antiparallel fibers that can be tilted out of the image plane and crossed in 2D or 3D configuration. The ultrastructural information about the fibrillar structures obtained from the polarimetric SHG imaging can be used to study, for example, changes in collagenous tissue due to tumor development, wound healing, fibrosis, and aging (28, 54–59).

## 6 Conclusion

SHG responses of chiral fibers with different configurations in the focal volume of nonlinear microscope have been analytically modelled and the outcomes compared with the results of PIPO measurements for tendon, cornea and articular cartilage tissues, and fibrillar TPPS_4_ aggregates. There are several fiber configurations in the focal volume that strongly influence the SHG response: (i) Parallel fibers dramatically increase and antiparallel fibers diminish the SHG intensity due to coherent summation of the signals generated from individual fibers. The R and C ratio values for parallel/antiparallel configurations are similar to the molecular ratio values of a single fiber; (ii) R and C ratios increase by tilting fibers out of the image plane for the molecular structures with R^*′*^ *<*3 (e.g. collagen). R ratio decreases with tilt angle *α* for structures with molecular R^*′*^ *>*3 (e.g. TPPS_4_ fibers), while C ratio increases with the tilt. The sign of C ratio depends on the tilt direction from the image plane (*α* or − *α*); (iii) The fiber crossing angle *σ* affects effective R and C ratios. R and C ratios increase with increasing *σ* for structures with molecular R^*′*^ *<*3, and R and C ratios decrease for structures with R^*′*^ *>*3. Both R and C ratios can be modified when a combination of tilting and crossing effects appear in the fibrillar organization of the tissue. The experimental results of PIPO SHG microscopy are consistent with the modeling results of digital nonlinear microscope presented in the previous paper (20).

This study contributes to the understanding and interpretation of imaging results of biological tissues obtained with polarimetric SHG microscopy. The results demonstrate that polarimetric SHG microscopy is a powerful imaging technique enabling to decipher 3D ultrastructural organization of fibrillar structures in biological specimens.

## Data availability

The datasets used during the current study are available from the corresponding author on reasonable request.

## Author Contributions

V.B. and M.A. developed the investigation plan of SHG with different samples. M.A and F.H measured the samples. M.A, V.B., and S.K. derived analytical expressions. L.K. and M.M. built the microscope. S.B., V.B., D.B., V.K., R.R., and M.K.A. prepared the samples. M.A. and V.B. interpreted the results and wrote the article. All authors contributed to the manuscript.

